# Positive and negative intra-molecular modulation in a dual-cassette RNA helicase

**DOI:** 10.1101/758698

**Authors:** Karen Vester, Karine F. Santos, Benno Kuropka, Christoph Weise, Markus C. Wahl

## Abstract

RNA helicase Brr2 is required for the activation of the spliceosome prior to the first catalytic step of splicing. Brr2 represents a distinct subgroup of Ski2-like nucleic acid helicases whose members comprise tandem helicase cassettes. Only the N-terminal cassette of Brr2 is an active ATPase and can unwind substrate RNAs. The C-terminal cassette represents a pseudo-enzyme that can stimulate RNA-related activities of the N-terminal cassette. However, the molecular mechanisms, by which the C-terminal cassette modulates the activities of the N-terminal unit remain elusive. Here, we show that N- and C-terminal cassettes adopt vastly different relative orientations in a crystal structure of Brr2 in complex with an activating domain of the spliceosomal Prp8 protein as compared to the crystal structure of isolated Brr2. Likewise, the cassettes occupy different relative positions and engage in different inter-cassette contacts during different stages of splicing. Engineered disulfide bridges that lock the cassettes in two different relative orientations have opposite effects on RNA-related activities of the N-terminal cassette compared to the unrestrained protein. Moreover, different relative positioning of the cassettes strongly influences ATP hydrolysis by the N-terminal cassette. Our results demonstrate that the inactive C-terminal cassette of Brr2 can exert both positive and negative influence on the active N-terminal helicase unit from a distance.

## INTRODUCTION

Nucleic acid-dependent nucleotide tri-phosphatases (NTPases) are present in all domains of life, as these enzymes are involved in virtually all aspects of gene expression and regulation (1). A subset of these NTPases acts on RNAs. Many RNA-dependent NTPases can function as RNA helicases *in vitro, i.e.* they unwind RNA duplexes in an NTP-dependent manner (2,3). *In vivo*, they might also perform other functions, such as RNA annealing (4,5), RNA clamping (6), displacement of RNA-bound proteins (7) or displacement of RNA-bound RNA-protein complexes (RNPs) (8,9).

To carry out these functions, RNA-dependent NTPases rely on a set of about twelve conserved sequence motifs that are involved in NTP binding, NTP hydrolysis, RNA binding and coupling of NTP binding/hydrolysis/product release to RNA/RNP transactions (10). Based on the presence and the degree of conservation of particular motifs, RNA-dependent NTPases have been assigned to several superfamilies (SFs) of nucleic acid-dependent NTPases, each of which contains several distinct families and subfamilies of enzymes (11). The vast majority of RNA-dependent NTPases belong to SF1 and SF2, whose members typically act as monomers (11). SF1/2 RNA-dependent NTPases all comprise a core of two RecA-like motor domains, which embody the NTP and RNA/RNP-related activities. Depending on the specific family, the proteins can contain additional domains, which are appended to or inserted into the RecA-like domains, as well as intrinsically unstructured terminal extensions (12). These helicase-associated domains and regions can modulate the activities of the enzymes in diverse ways (12).

Several families of RNA-dependent NTPases also contain a conserved set of helicase-associated domains, which, together with the RecA-like core domains, form functional NTPase/helicase units or cassettes. For example, members of the Ski2-like subfamily of SF2 harbor helicase cassettes, in which the two RecA-like domains are followed by a winged-helix (WH), a helical bundle (HB) and a helix-loop-helix (HLH) domain, and most members comprise an additional C-terminal immunoglobulin-like (IG) domain. A small subgroup of Ski2-like enzymes contains duplicated helicase cassettes arranged in tandem. The genomes of yeast and human each encode two such dual-cassette enzymes, *i.e.* Slh1/Rqt2 and Brr2 in yeast and ASCC3 and Brr2 in human. ASCC3 is a subunit of the activating signal cointegrator complex, which was originally identified as a coactivator of nuclear receptors (13) and later found to be involved in down-tuning of cellular anti-viral responses (14), regulation of myogenic differentiation (15,16) and DNA dealkylation repair (17,18). Slh1/Rqt2 may constitute the yeast ortholog of human ASCC3 (19).

Brr2 is an RNA helicase involved in precursor messenger RNA (pre-mRNA) splicing and is presently the best investigated representative of the dual-cassette Ski2-like helicases. Pre-mRNA splicing is carried out by a large and dynamic RNP molecular machine, the spliceosome (20). The spliceosome assembles *de novo* on each pre-mRNA substrate in a stepwise manner, is then catalytically activated, carries out two transesterification reactions that lead to intron excision and exon ligation and is finally disassembled in an ordered manner (20). The transitions between the assembly, activation, catalysis and disassembly stages are accompanied by profound compositional and conformational remodeling of the spliceosome, which is promoted by at least eight conserved, spliceosome-associated RNA-dependent NTPases/RNA helicases. Brr2 is involved in the conversion of a pre-catalytic spliceosomal B complex to an activated B^act^ complex. In that process, Brr2 unwinds the initially base-paired spliceosomal U4 and U6 small nuclear (sn) RNAs (21), enabling U6 to adopt a new structure, engage in new interactions and form part of the spliceosome’s active site (22). Regulation of the spliceosomal RNA helicases has been implicated in splicing fidelity (23) and in the regulation of alternative splicing (24), *i.e.* the inclusion of different combinations of coding regions in mature mRNAs, by kinetic proofreading mechanisms and by funneling spliceosomal assembly intermediates into discard pathways (25). Thus, regulation of the activities of the spliceosomal helicases may have a major impact on gene expression and regulation, in particular in higher eukaryotes where alternative splicing is pervasive (26).

A number of specific regulatory principles have been delineated for Brr2 (27). For example, besides the two helicase cassettes, Brr2 comprises a large N-terminal region of about 400 residues that can fold back onto the helicase cassettes and auto-inhibit the enzyme by substrate competition and conformational clamping (28). In addition, a C-terminal Jab1 domain of the spliceosomal master regulator, Prp8, can either inhibit or activate Brr2, depending on whether or not a C-terminal tail of the domain is inserted into the helicase’s RNA-binding tunnel (29-31). Other proteins binding to Brr2 can also modulate its activities *in vitro* (32,33).

Only the N-terminal cassette (NC) of Brr2 is an active NTPase/RNA helicase, while the C-terminal cassette (CC) is considered a pseudo-enzyme (13,34). Nevertheless, the CC is essential for yeast viability (35), can still bind but does not hydrolyze ATP (34) and serves as an interaction platform for a number of other splicing factors (36,37), some of which have been shown to modulate Brr2 activity by presently unknown mechanisms (32,33). Indeed, in human (h) Brr2, the CC itself can activate the NC helicase (34). Presently, the regulatory potential of the Brr2 CC has not been thoroughly explored, and the molecular mechanisms underlying Brr2 regulation *via* its CC are poorly understood.

Here, we present direct evidence for the ability of the two hBrr2 cassettes to occupy vastly different relative positions and engage in different inter-cassette interactions in isolation and during different stages of splicing. Upon fixing two such relative arrangements by disulfide bridges, we observed opposite effects on RNA/ATP-related activities of the NC. Our results show that the CC can function as an intra-molecular cofactor of the NC helicase, which, depending on its relative position and inter-cassette interactions, can exert positive or negative regulatory effects.

## RESULTS

### The hBrr2 helicase cassettes can adopt diverse relative positions and orientations

We had previously observed slight changes in relative cassette orientations in the crystal structures of a dual cassette-fragment of hBrr2 (residues 394-2129; here referred to as hBrr2 truncation 1, hBrr2^T1^; Fig. 1A), in isolation (34) and in complex with the hPrp8 Jab1 domain, bearing a C-terminal tail in a hBrr2-inhibitory configuration (29). To further explore the inter-cassette flexibility in hBrr2, we determined the crystal structure of hBrr2^T1^ in complex with a hPrp8 Jab1 domain lacking the inhibitory tail (residues 2064-2320; hJab1^ΔCtail^), which we had previously shown to be a strong activator of the hBrr2 helicase (29). Crystals of the hBrr2^T1^-hJab1^ΔCtail^ complex diffracted to 2.4 Å resolution and the structure was solved by molecular replacement using the structure coordinates of isolated hBrr2^T1^ (PDB ID 4F91) (34) and of the tail-deleted hPrp8 Jab1 domain (PDB ID 4KIT) (29) (Table 1). In the hBrr2^T1^-hJab1^ΔCtail^ structure, hBrr2^T1^ adopted a vastly different relative cassette orientation compared to hBrr2^T1^ alone or compared to hBrr2^T1^ within a hBrr2^T1^-hJab1 complex (Fig. 1B,C). Relative to isolated hBrr2^T1^, the apparent movement of the CC relative to the NC in the hBrr2^T1^-hJab1^ΔCtail^ structure is described by a rotation of 21.8 ° around the linker between NC and CC (apparent rotation from the hBrr2^T1^ structure to the hBrr2^T1^-hJab1^ΔCtail^ structure defined as positive; Fig. 2, first panel; rot+). These observations show that the relative positions of the two cassettes in hBrr2 can cover a much larger spectrum than previously realized.

**Table 1.** Crystallographic data^a^.

| Dataset | hBrr2<sup>T1</sup>-<br>hJab1<sup>ΔCtail</sup> | hBrr2<sup>T1-M641C/A1582C</sup> | hBrr2<sup>T1-D534C/N1866C</sup>-<br>hJab1<sup>ΔCtail</sup> |
| --- | --- | --- | --- |
| **Data Collection** |  |  |  |
| Wavelength [Å] | 0.91842 | 0.91840 | 0.91677 |
| Temperature [K] | 100 | 100 | 100 |
| Space group | P2<sub>1</sub>2<sub>1</sub>2<sub>1</sub> | C2 | P2<sub>1</sub>2<sub>1</sub>2<sub>1</sub> |
| Unit cell parameters |  |  |  |
| Axes [Å] | 99.5, 118.7, 187.8 | 145.6, 152.5, 142.5 | 99.2, 119.1, 187.9 |
| Angles [°] | 90.0, 90.0, 90.0 | 90.0, 120.7, 90.0 | 90.0, 90.0, 90.0 |
| Resolution [Å] | 50.0-2.4 (2.53-2.39) | 50.0-3.2 (3.36-3.17) | 50.0-2.6 (2.76-2.60) |
| Reflections |  |  |  |
| Unique | 88725 (14026) | 44087 (6492) | 68538 (10962) |
| Completeness [%] | 99.8 (99.0) | 97.0 (89.3) | 98.9 (99.2) |
| Redundancy/ multiplicity | 8.9 (8.4) | 3.4 (3.4) | 5.7 (5.6) |
| I/σ(I) | 11.3 (1.7) | 8.7 (1.0) | 9.6 (1.3) |
| R<sub>meas</sub>(I) [%]<sup>b</sup> | 14.4 (144.6) | 12.1 (131.6) | 15.9 (138.6) |
| CC<sub>1/2</sub> [%]<sup>c</sup> | 99.7 (59.4) | 99.8 (64.1) | 99.7 (47.4) |
| **Refinement** |  |  |  |
| Resolution [Å] | 48.1 - 2.4 (2.48 - 2.39) | 48.4 - 3.2 (3.33 - 3.17) | 47.9 - 2.6 (2.69 - 2.59) |
| Reflections |  |  |  |
| Number | 88212 (8654) | 44032 (5199) | 68531 (6757) |
| Completeness [%] | 99.73 (98.84) | 97.2 (88.7) | 98.85 (98.77) |
| Test set | 2100 (206) | 2094 (247) | 2099 (207) |
| R<sub>work</sub><sup>d</sup> | 17.9 (25.8) | 26.4 (44.7) | 19.6 (31.4) |
| R<sub>free</sub><sup>e</sup> | 25.3 (33.9) | 31.8 (52.3) | 27.9 (39.8) |
| Contents of A.U.<sup>f</sup> |  |  |  |
| Protein atoms | 16866 | 13858 | 15823 |
| Ligand atoms | 10 | - | - |
| Water oxygens | 656 | 20 | 102 |
| Mean B factors [Å<sup>2</sup>] |  |  |  |
| Wilson | 56.6 | 94.0 | 58.2 |
| Protein | 59.6 | 118.9 | 69.2 |
| Ligands | 121.1 | - | - |
| Water oxygens | 49.2 | 107.2 | 49.7 |
| Ramachandran plot<sup>g</sup> |  |  |  |
| Favored [%] | 95.00 | 98.90 | 93.04 |
| Outliers [%] | 0.76 | 1.10 | 1.07 |
| Rmsd<sup>h</sup> |  |  |  |
| Bond lengths [Å] | 0.009 | 0.002 | 0.006 |
| Bond angles [°] | 1.10 | 0.54 | 0.94 |
| PDB ID | 6S8Q | 6S8O | 6S9I |

<sup>a</sup> Values in parentheses refer to the highest resolution shells.<sup>b</sup> $R_{\text{meas}}(I) = \sum_h [N/(N-1)]^{1/2} \sum_i |I_{ih} - \langle I_h \rangle| / \sum_h \sum_i I_{ih}$ , in which $\langle I_h \rangle$ is the mean intensity of symmetry-equivalent reflections $h$ , $I_{ih}$ is the intensity of a particular observation of $h$ and $N$ is the number of redundant observations of reflection $h$ .
- <sup>c</sup> $CC_{1/2} = (\langle I^2 \rangle - \langle I \rangle^2) / (\langle I^2 \rangle - \langle I \rangle^2) + \sigma_e^2$ , in which $\sigma_e^2$ is the mean error within a half-dataset (74). <sup>d</sup> $R_{work} = \sum_h |F_o - F_c| / \sum F_o$ (working set, no $\sigma$ cut-off applied). <sup>e</sup> $R_{free}$ is the same as $R_{work}$ , but calculated on the test set of reflections excluded from refinement. <sup>f</sup> A.U. – asymmetric unit. <sup>g</sup> Calculated with Phenix. <sup>h</sup> Rmsd - root-mean-square deviation from target geometry.

**Figure 1.**
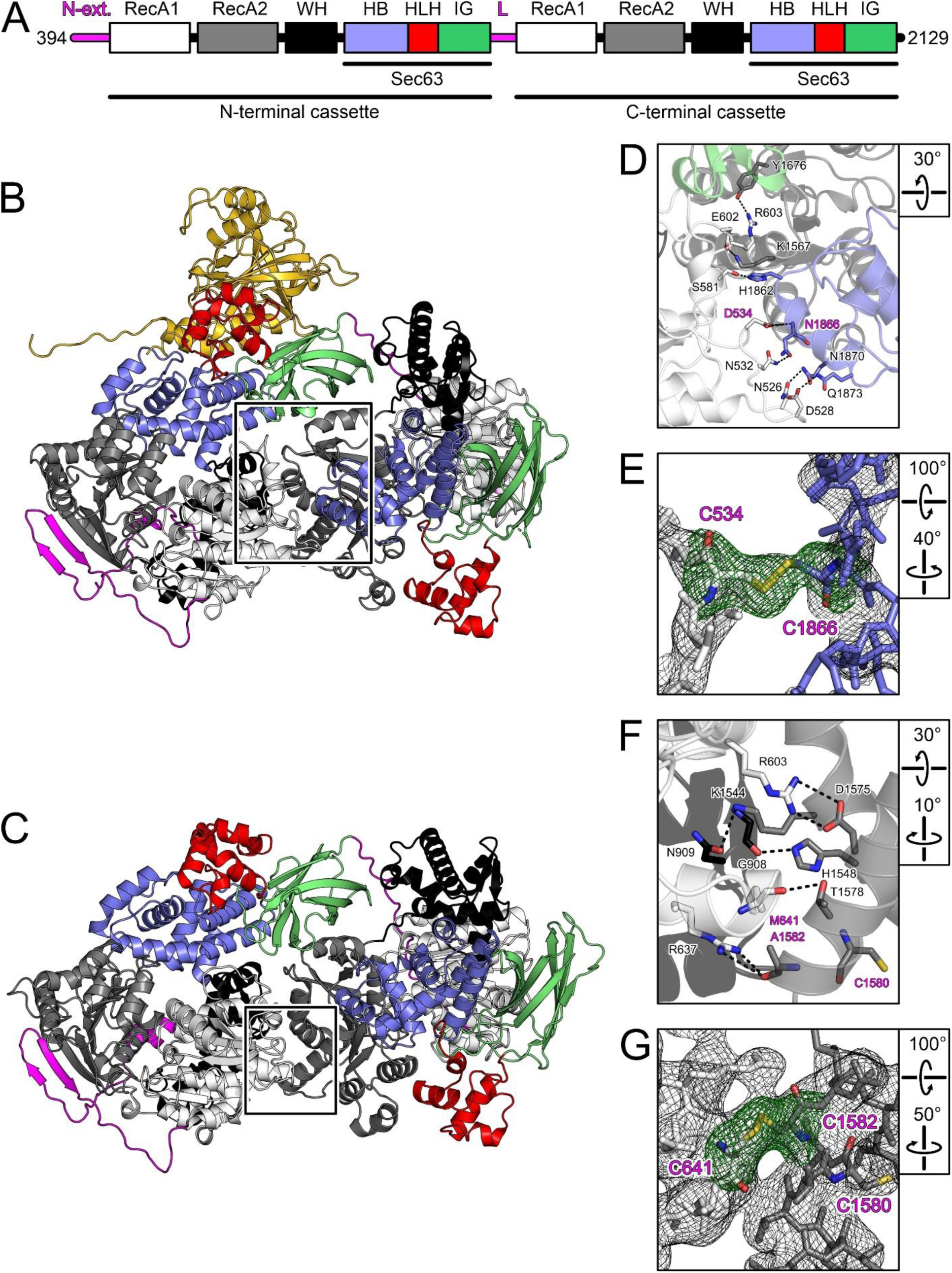
Structures of hBrr2^T1^ and introduction of disulfide bridges. (A) Domain structure of hBrr2^T1^. N-ext., N-terminal extension; RecA, RecA-like domains, WH, winged-helix domains; HB, helical bundle domains; HLH, helix-loop-helix domains; IG, immunoglobulin-like domains; L, inter-cassette linker; Sec63, Sec63 homology units. (B) Structure of the hBrr2^T1^-hJab1^ΔCtail^ complex. hBrr2^T1^ domains, colored as in (A); hJab1^ΔCtail^, gold. (C) Structure of the hBrr2^T1^ (34). hBrr2^T1^ domains, colored as in (A). (D) Zoom into the inter-cassette interface in the hBrr2^T1^-hJab1^ΔCtail^ complex (region boxed in (B)). In this and the following figure panels: Interacting residues shown as sticks. Carbon, as the respective protein region; nitrogen, blue; oxygen, red. Dashed lines, hydrogen bonds or salt bridges. Residues mutated to cysteines to introduce a disulfide bridge are labeled in magenta. Rotation symbols represent the view relative to panels (B) and (C). (E) 2Fo-Fc electron density (gray mesh; 1 σ level) and Fo-Fc “omit” electron density (green mesh; 3 σ level) around the engineered disulfide bridge in hBrr2^T1-D534C/N1866C^ (hBrr2^T1-SS-rot+^). hBrr2^T1-D534C/N1866C^ was crystallized in complex with hJab1^ΔCtail^. Mutated residues labeled in magenta. (F) Zoom into the inter-cassette interface in isolated hBrr2^T1^ (region boxed in (C)). (G) 2Fo-Fc electron density (gray mesh; 1 σ level) and Fo-Fc “omit” electron density (green mesh; 3 σ level) around the engineered disulfide bridge in hBrr2^T1-M641C/A1582C^ (hBrr2^T1-SS-linear^). hBrr2^T1-M641C/A1582C^ was crystallized in isolation. Mutated residues and a neighboring cysteine are labeled in magenta.

**Figure 2.**
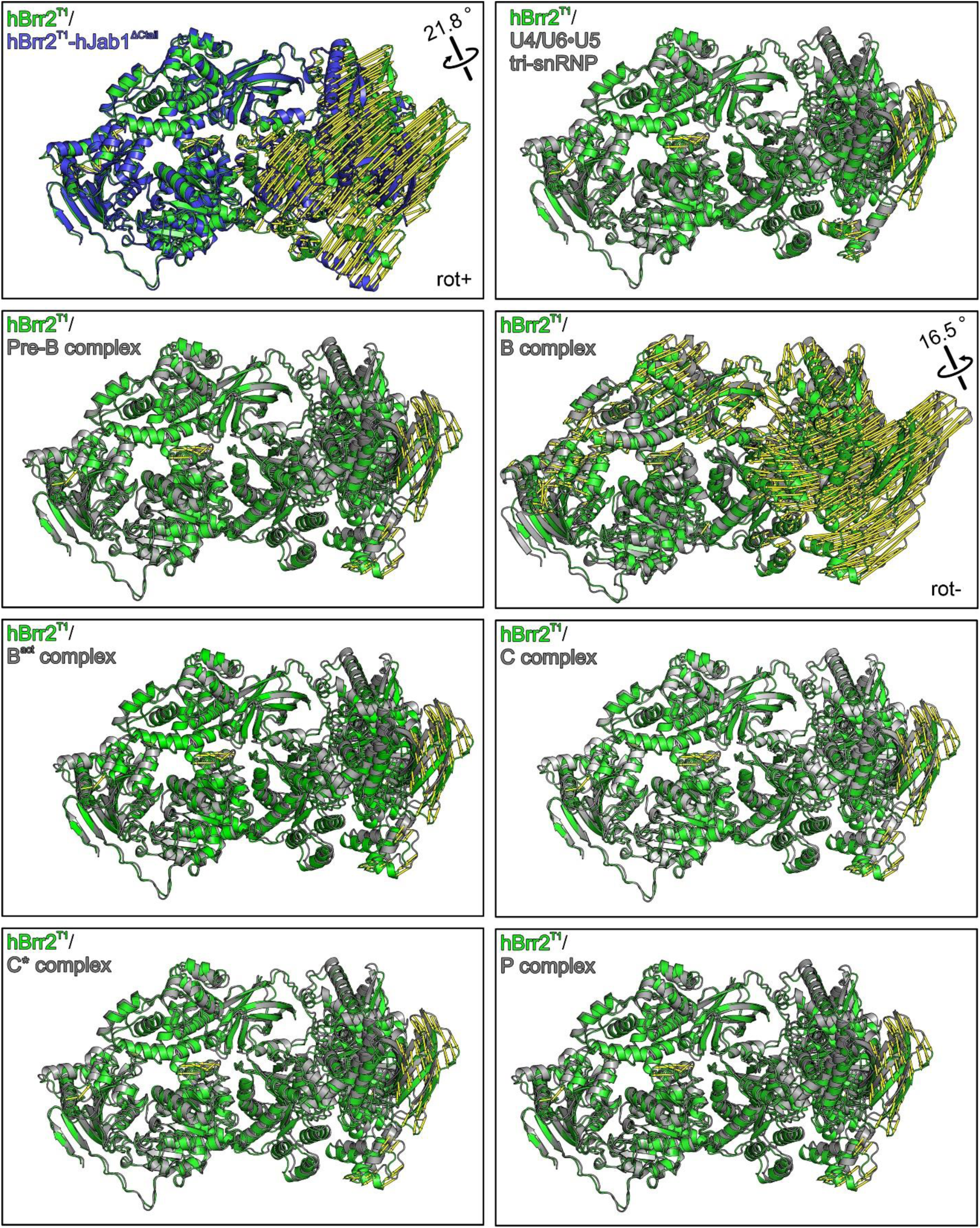
Structural comparisons. Crystal structure of hBrr2^T1^ (green) (34) superimposed on the hBrr2^T1^-hJab1^ΔCtail^ hJab1^ΔCtail^ structure (blue) and on the hBrr2 subunits in cryoEM structures of the U4/U6•U5 tri-snRNP (39), spliceosomal pre-B (40), B (38), B^act^ (42), C (43), C* (41) and P complexes (44). Alignments were based on NC residues 1-1288. Yellow “mode” vectors indicate structural differences as distances after alignment (displacements of common Cα positions of the compared structures to the isolated hBrr2^T1^ reference structure). rot+, apparent rotation of CC in the hBrr2^T1^ structure to the CC in the hBrr2^T1^-hJab1^ΔCtail^ structure defined as positive; rot-, opposite apparent rotation sense for the CC seen in all structures of spliceosomal complexes, most prominently in the B complex.

The different relative positioning of the cassettes in the hBrr2^T1^-hJab1^ΔCtail^ complex crystals may reflect conformational changes induced by the activatory hJab1^ΔCtail^ or may be the consequence of crystal packing effects, as the hBrr2^T1^-hJab1^ΔCtail^ complex crystallized in a different space group than hBrr2^T1^ or hBrr2^T1^-hJab1 (Table 1). To investigate whether hBrr2 also can undergo inter-cassette conformational changes in the framework of spliceosomal complexes, we compared hBrr2^T1^ crystal structures to the conformations of hBrr2 subunits extracted from recently published cryo-electron microscopy (cryoEM) structures of human spliceosomes (38-44). Structural alignments *via* the NCs revealed that the CCs can adopt different relative orientations in different functional contexts (Fig. 2; Table 2). A similar picture emerged when we compared the conformations of Brr2 subunits extracted from cryoEM structures of yeast spliceosomal complexes (45-50) (Table 2). hBrr2 conformations of all cryoEM structures aligned more closely to the conformation of isolated hBrr2^T1^ than to the conformation of hBrr2^T1^ in the crystal structure of the hBrr2^T1^-hJab1^ΔCtail^ complex (Table 2). Moreover, compared to isolated hBrr2^T1^, all hBrr2 conformations in spliceosomal cryoEM structures exhibited rotations of the CC relative to the NC in the opposite sense compared to hBrr2^T1^ in the hBrr2^T1^-hJab1^ΔCtail^ complex, with Brr2 in the B complex exhibiting the largest rotation of 16.5 ° (Fig. 2; rot-). Brr2 resides in a peripheral, less well-resolved region in most spliceosomal cryoEM structures, which may obscure the full extent of Brr2 conformational changes in the imaged contexts, and not all assembly, activation and catalysis intermediates of a splicing cycle have so far been structurally analyzed. Thus, the conformational spectrum explored by Brr2 during a splicing cycle might exceed the conformational flexibility portrayed by the presently available spliceosome structures. Irrespectively, these analyses reveal that Brr2 apparently undergoes inter-cassette conformational changes during a splicing cycle.

**Table 2.** Structural comparisons.

| Structure | PDB ID | Rmsd<sup>a</sup> hBrr2<sup>T1</sup> [Å<sup>2</sup>] | Rmsd hBrr2<sup>T1</sup>-Jab1<sup>ΔCtail</sup> [Å<sup>2</sup>] |
| --- | --- | --- | --- |
| human U4/U6•U5 tri-snRNP | 3JCR | 1.5 | 5.6 |
| human B complex | 5O9Z | 2.7 | 9.6 |
| human B<sub>act</sub> complex | 5Z57 | 1.5 | 5.7 |
| human C complex | 5YZG | 1.5 | 5.7 |
| human C* complex | 5XJC | 1.5 | 5.7 |
| yeast U4/U6•U5 tri-snRNP | 3JCM | 5.6 | 9.3 |
| yeast U4/U6•U5 tri-snRNP | 5GAO | 5.9 | 9.6 |
| yeast B complex | 5NRL | 5.9 | 9.7 |
| yeast B<sub>act</sub> complex | 5GM6 | 4.4 | 6.6 |
| yeast B<sub>act</sub> complex | 5LQW | 4.1 | 5.7 |
| yeast C complex | 5LJ5 | 3.0 | 6.5 |

<sup>h</sup> Rmsd - root-mean-square deviation from target geometry.

### Alternative hBrr2 conformations can be stabilized via engineered disulfide bridges

The hBrr2^T1^ conformations in the isolated and hBrr2^T1^-hJab1^ΔCtail^ crystal structures exhibit major differences in the contacts of the CC to the NC at the interface between the cassettes (Fig. 1D,F). Whereas the conformation of isolated hBrr2^T1^ encompasses several salt bridges and hydrogen bonds between the N-terminal RecA1/WH domains and the C-terminal RecA2 domain (Fig. 1F), the cassette interface in the hBrr2^T1^-hJab1^ΔCtail^ structure comprises fewer and longer-distance polar and hydrophobic interactions between the N-terminal RecA1 and the C-terminal RecA2/HB domains (Fig. 1D). In the following, we refer to the relative cassette arrangement in isolated hBrr2^T1^ and hBrr2^T1^-hJab1^ΔCtail^ complex as the linear and positively rotated (rot+) conformations.

Previously, we had shown that exchanges of residues, which foster inter-cassette contacts in the linear conformation of isolated hBrr2^T1^, or in the extended linker region connecting the two cassettes, led to alterations in the NC helicase activity (34). We therefore set out to test whether the two conformations represented by isolated hBrr2^T1^ and hBrr2^T1^-hJab1^ΔCtail^ crystal structures would correlate with differences in the helicase activity of hBrr2. To this end, we selected pairs of NC/CC residues that resided in suitable relative positions in the two structures to lead to the formation of disulfide bridges upon joint exchange for cysteine residues, thereby stabilizing the linear or the rot+ conformation (Fig. 1D,F). Cysteine residues were introduced at the selected positions (M641C/A1582C for the linear conformation of isolated hBrr2^T1^; D534C/N1866C for the rot+ conformation of hBrr2^T1^-hJab1^ΔCtail^) by site-directed mutagenesis, the corresponding hBrr2^T1^ variants were purified (but omitting addition of DTT in the final gel filtration step; Fig. S1) and crystallized alone (hBrr2^T1-M641C/A1582C^; in the following referred to as hBrr2^T1-SS-linear^) or in complex with hJab1^ΔCtail^ (hBrr2^T1-D534C/N1866C^; in the following referred to as hBrr2^T1-SS-rot+^).

Crystal structure analyses unequivocally revealed the presence of the intended disulfide bridges (Fig. 1E,G). Moreover, structural alignments showed that the disulfide-bridged hBrr2^T1^ variants adopted essentially the same conformations as the WT variant in the two different structural contexts (isolated hBrr2^T1^/hBrr2^T1-SS-linear^, root-mean-square deviation [rmsd] of 0.519 Å for 1,485 matching pairs of hBrr2^T1^ Cα atoms; hBrr2^T1^/hBrr2^T1-SS-rot+^-Jab1^ΔCtail^, rmsd of 0.278 Å for 1,523 matching pairs of hBrr2^T1^ Cα atoms).

To test whether the intended disulfide bridges also formed in solution prior to crystallization, we compared the fold stabilities of WT and cysteine-derivatized hBrr2^T1^ using differential scanning fluorimetry (DSF) in two different buffers and in the presence or absence of the reducing agent DTT (Fig. 3A,B). Not surprisingly, the buffer composition *per se* did have an influence on the melting temperatures of the proteins. However, in both buffer systems, the presence or absence of DTT influenced the melting temperatures of the cysteine variants, but not those of the WT protein, in the same way and in complete agreement with the formation of the intended disulfide bridges under non-reducing conditions. *I.e.*, in the absence of DTT, melting temperatures of both cysteine-variants were increased by 2-3 ° C in the two buffers compared to those of the WT protein (Fig. 3A,B, columns 3 and 5). In contrast, in the presence of DTT, the melting temperatures of both cysteine-variants were indistinguishable and essentially identical to those of the WT protein in DTT-containing buffers (Fig. 3A,B, columns 4 and 6).

**Figure 3.**
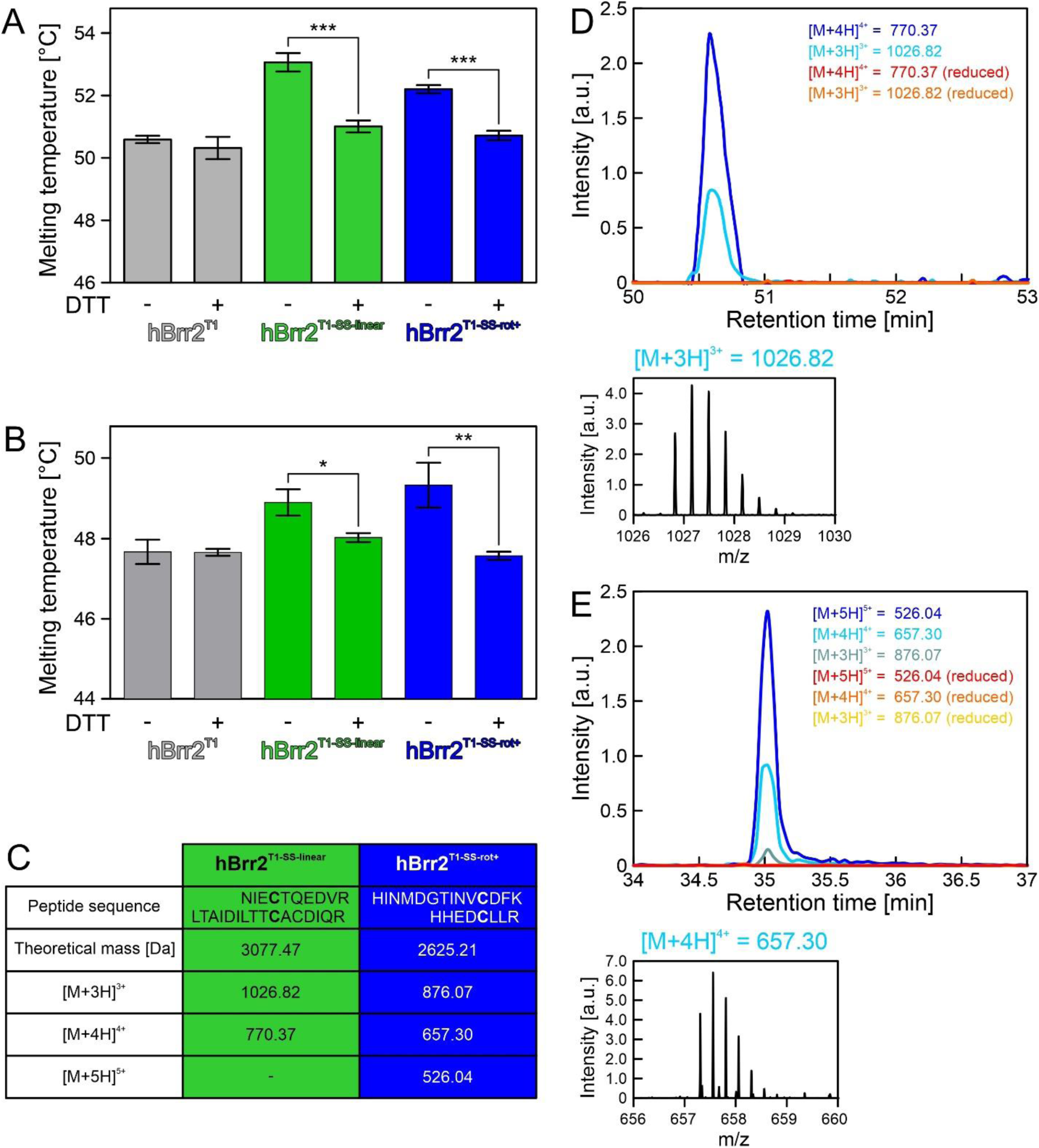
Analysis of the hBrr2^T1^ bearing engineered disulfide bridges. (A) DSF-derived melting temperatures for hBrr2^T1^ (gray), hBrr2^T1-SS-linear^ (green) and hBrr2^T1-SS-rot+^ (blue) with and without addition of the reducing agent, DTT, in SEC buffer. Values represent means ± SD of at least three independent experiments. ***, p ≤ 0.001. (B) DSF-derived melting temperatures for hBrr2^T1^ (gray), hBrr2^T1-SS-linear^ (green) and hBrr2^T1-SS-rot+^ (blue) with and without addition of the reducing agent, DTT, in 40 mM TRIS-HCl, pH 7.5, 50 mM NaCl, 0.5 mM MgCl2. Values represent means ± SD of at least three independent experiments. *, p ≤ 0.05; **, p ≤ 0.01. (C) Mass-to-charge ratios (m/z) of disulfide-bridged peptides identified in LC-MS analyses. (D) Extracted ion chromatograms for disulfide-bridge containing peptides of hBrr2^T1-SS-linear^ present in the non-reduced/oxidized samples and missing in the reduced samples. One exemplary MS1 spectrum of the triply charged bridged peptide is shown. (E) Extracted ion chromatograms for disulfide-bridge containing peptides of hBrr2^T1-SS-rot+^ present in the non-reduced/oxidized samples and missing in the reduced samples. One exemplary MS1 spectrum of the quadruply charged bridged peptide is shown.

To further test formation of the engineered disulfide bridges in solution, we conducted mass spectrometric (MS) analyses (Fig. 3C-E). Free thiol groups in hBrr2^T1-SS-linear^ and hBrr2^T1-SS-rot+^ were modified with N-ethylmaleimide (NEM) before or after reduction with DTT, and subsequently proteins were digested with LysC and trypsin. MS1 signals corresponding to the expected disulfide-bridged peptides (638-NIECTQEDVR-647/1571-LTAIDILTTCACDIQR-1586 for hBrr2^T1-SS-linear^; 524-HINMDGTINVCDFK-537/1862-HHEDCLLR-1869 for hBrr2^T1-SS-rot+^) were identified in the non-reduced samples (Fig. 3C). For hBrr2^T1-SS-linear^, two different charge states of the bridged peptide were seen, while three different charge states were detected for the bridged peptide of hBrr2^T1-SS-rot+^ (Fig. 3C; examples for MS1 peaks are shown in Fig. 3D,E). Extracted-ion chromatograms of the reduced samples lacked signals of the disulfide-bridged peptides (Fig. 3D,E), confirming that the identified peaks in the non-reduced samples originated from disulfide-bridged peptides. The identities of the bridged peptides were further confirmed by MS2 spectra of material in peaks m/z 770.4 (hBrr2^T1-SS-linear^) and m/z 657.3 (hBrr2^T1-SS-rot+^; Fig. S2). The latter analysis revealed that the m/z 770.4 peak that originated from hBrr2^T1-SS-linear^ corresponded to 638-NIECTQEDVR-647/1571-LTAIDILTTCACDIQR-1586 with a disulfide bridge between C641 and C1580 rather than the engineered C1582. Therefore, either a C641/C1580 disulfide bridge formed in hBrr2^T1-SS-linear^ in solution while a C641/C1582 disulfide bridge was favored in the crystal (Fig. 1G), or disulfide shuffling between the neighboring C1580 and C1582 took place during sample preparation forMS. Irrespectively, both disulfide-bridged configurations would stabilize hBrr2^T1-SS-linear^ in the linear conformation seen in isolated WT hBrr2^T1^.

To estimate the fraction of bridged and unbridged protein molecules, the intensities of the corresponding unbridged peptides were compared between the non-reduced and reduced samples. This analysis indicated a high degree of disulfide bridge formation in the hBrr2^T1-SS-linear^ variant, but a lower degree of disulfide bridge formation in the hBrr2^T1-SS-rot+^ variant.

Together, the above analyses suggest that the engineered variants indeed form the intended disulfide bridges in solution, albeit not at 100 %, and that the replacements of the selected residues in the variants with cysteine or the presence of DTT *per se* have no effect on the fold stabilities. They therefore also indicate that the linear and rot+ conformations observed in hBrr2^T1^ and hBrr2^T1^-hJab1^ΔCtail^ crystals, respectively, are also adopted by hBrr2^T1^ in solution. hBrr2^T1^ in solution apparently populates the two conformations to different extents, with a higher fraction of hBrr2^T1^ molecules in the linear conformation and a lower fraction in the rot+ conformation.

### Restraining the cassettes in selected conformations influences Brr2 helicase activity

To test the consequences of arresting hBrr2^T1^ in two different inter-cassette conformations, we monitored the effects of hBrr2^T1-SS-linear^ and hBrr2^T1-SS-rot+^ in gel-based, radioactive U4/U6 unwinding assays. In comparison to WT hBrr2^T1^, hBrr2^T1-SS-linear^ exhibited a significantly higher unwinding rate (1.645 ± 0.049 s^-1^ *vs.* 0.941 ± 0.040 s^-1^), whereas hBrr2^T1-SS-rot+^ showed a significantly lower activity (0.536 ± 0.012 s^-1^; Fig. 4; Table 3). Notably, upon inclusion of DTT in the unwinding buffer, the activity of hBrr2^T1-SS-linear^ decreased while the activity of hBrr2^T1-SS-rot+^ increased, both approaching the unwinding rates of the WT variant under the same conditions (Fig. 4; Table 3). These trends were significant as indicated by non-overlapping 95 % confidence intervals compared to the values without reducing agent (Table 3). Thus, the observed effects clearly depend on the presence of the engineered disulfide bridges. Given that engineered disulfide bridges are not formed at 100 % in solution (see above), the unwinding assays underestimate the effects of completely locking hBrr2^T1^ in either linear or rot+ conformation, in particular for the latter situation. These findings indicate that hBrr2 is more active in unwinding a biologically relevant RNA substrate when it is restrained in the conformation represented by the isolated hBrr2^T1^ crystal structure, while restraining hBrr2^T1^ in the conformation seen in the hBrr2^T1^-hJab1^ΔCtail^ complex structure leads to decreased activity.

**Table 3.**
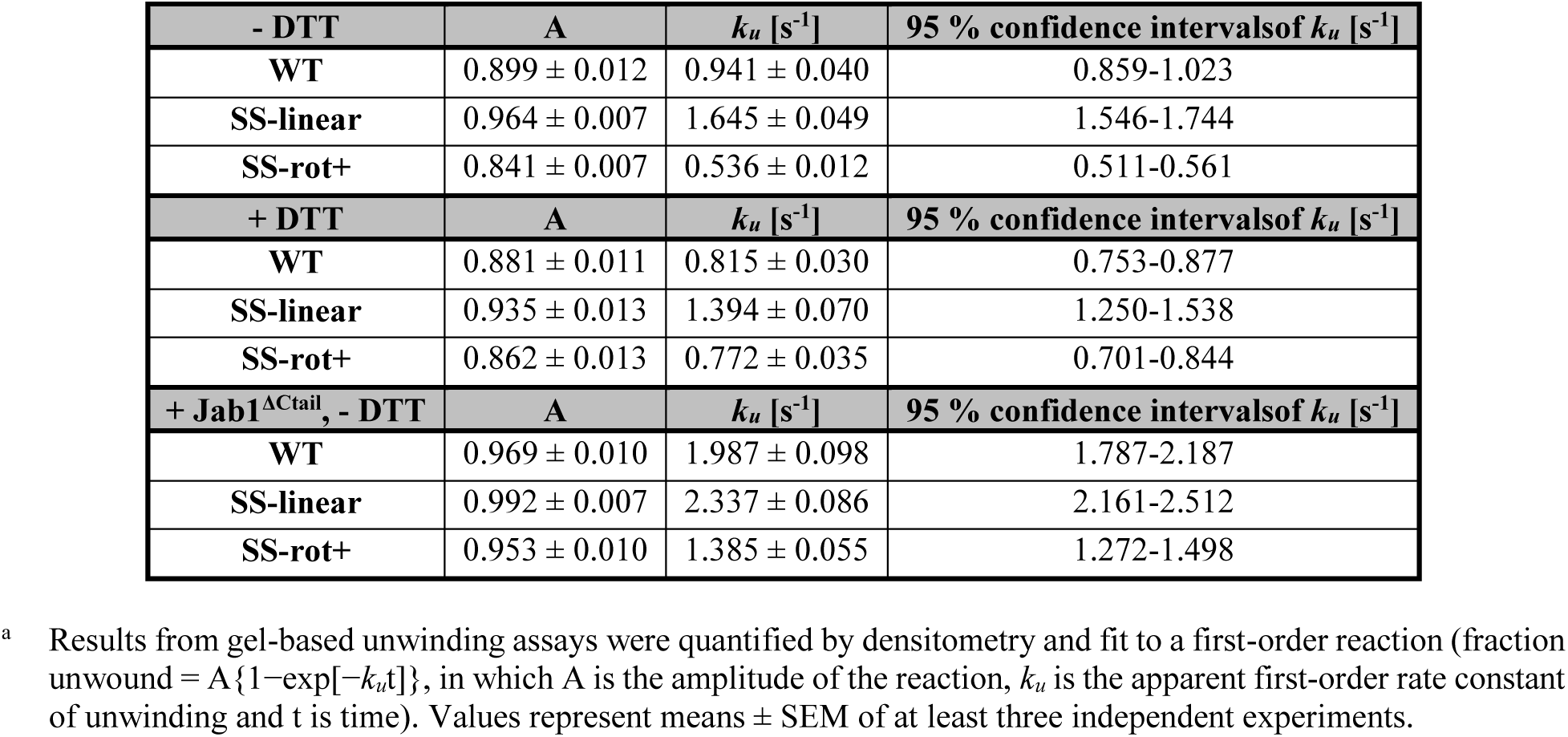
Amplitudes and rate constants of U4/U6 unwinding^a^.

| - DTT | A | $k_u$ [s<sup>-1</sup>] | 95 % confidence interval of $k_u$ [s<sup>-1</sup>] |
| --- | --- | --- | --- |
| WT | 0.899 ± 0.012 | 0.941 ± 0.040 | 0.859-1.023 |
| SS-linear | 0.964 ± 0.007 | 1.645 ± 0.049 | 1.546-1.744 |
| SS-rot+ | 0.841 ± 0.007 | 0.536 ± 0.012 | 0.511-0.561 |
| + DTT | A | $k_u$ [s<sup>-1</sup>] | 95 % confidence interval of $k_u$ [s<sup>-1</sup>] |
| WT | 0.881 ± 0.011 | 0.815 ± 0.030 | 0.753-0.877 |
| SS-linear | 0.935 ± 0.013 | 1.394 ± 0.070 | 1.250-1.538 |
| SS-rot+ | 0.862 ± 0.013 | 0.772 ± 0.035 | 0.701-0.844 |
| + Jab1<sup>ΔCtail</sup>, - DTT | A | $k_u$ [s<sup>-1</sup>] | 95 % confidence interval of $k_u$ [s<sup>-1</sup>] |
| WT | 0.969 ± 0.010 | 1.987 ± 0.098 | 1.787-2.187 |
| SS-linear | 0.992 ± 0.007 | 2.337 ± 0.086 | 2.161-2.512 |
| SS-rot+ | 0.953 ± 0.010 | 1.385 ± 0.055 | 1.272-1.498 |

<sup>a</sup> Results from gel-based unwinding assays were quantified by densitometry and fit to a first-order reaction (fraction unwound = $A \{1 - \exp[-k_u t]\}$ , in which A is the amplitude of the reaction, $k_u$ is the apparent first-order rate constant of unwinding and t is time). Values represent means ± SEM of at least three independent experiments.

**Figure 4.**
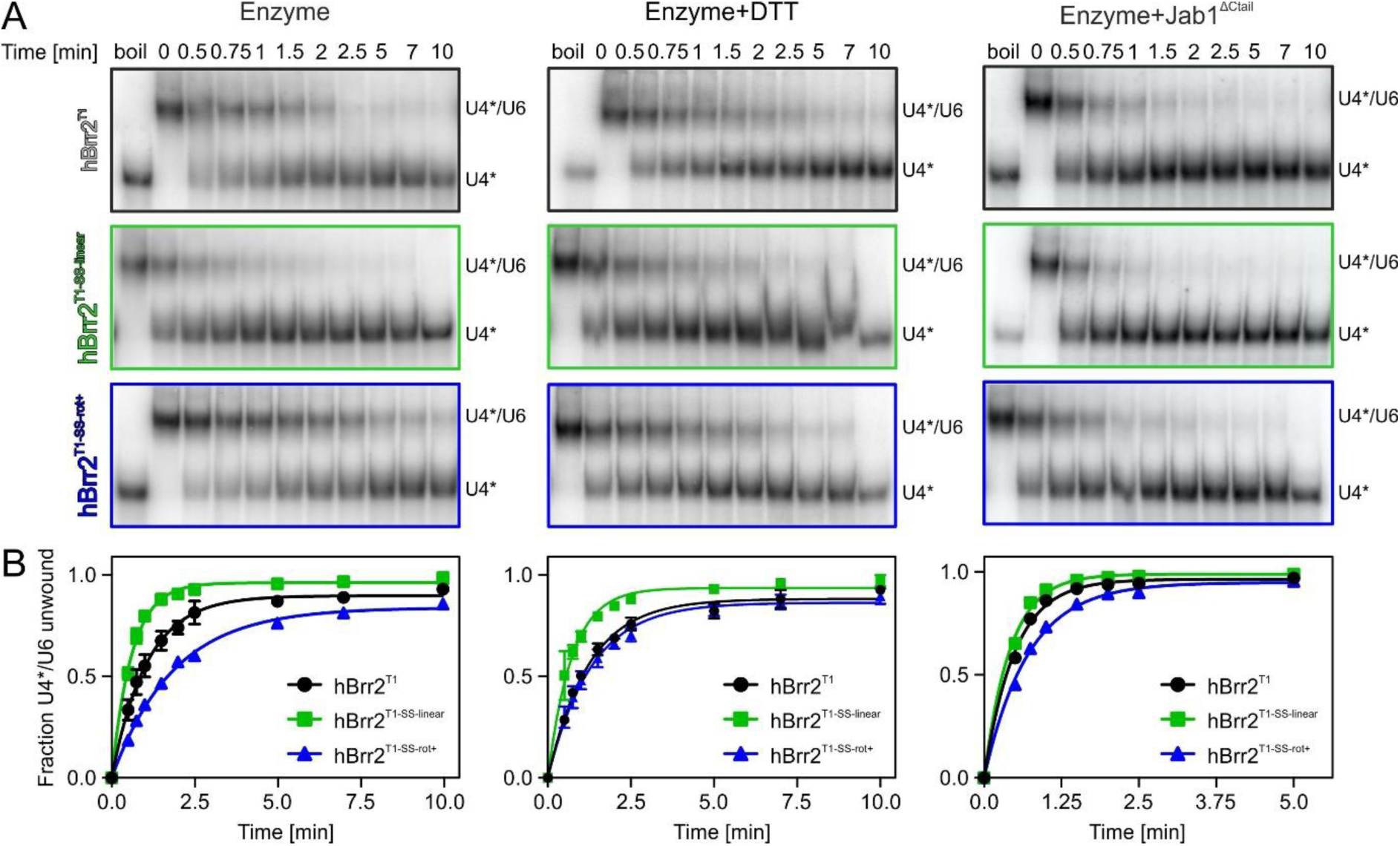
Helicase activities. (A) Exemplary gels monitoring unwinding of U4*/U6 duplexes by the hBrr2T1 variants indicated on the left. Left: Enzyme alone, untreated. Middle: Enzyme alone, treated with reducing agent. Right: Enzyme in complex with hJab1^Δctail^. U4*, radioactively labeled U4 snRNA. (B) Quantification of the data shown in (A). hBrr2^T1^, black; hBrr2^T1-SS-linear^, green; hBrr2^T1-SS-rot+^, blue. Data points represent means ± SD of at least three independent experiments.

### The hPrp8 Jab1 domain does not alter the trends in activity modulation upon crosslinking

To test whether the effects on hBrr2^T1^ helicase activity prevail in the presence of hJab1^ΔCtail^, we repeated the unwinding assays in the presence of the activator. As expected, hJab1^ΔCtail^ increased the unwinding rates of all hBrr2^T1^ constructs, consistent with previous studies (29) (Fig. 4; Table 3). These results show that hJab1^ΔCtail^ can still bind to hBrr2^T1^ when restrained in the linear conformation seen in the isolated hBrr2^T1^ crystal structure (hBrr2^T1-SS-linear^), consistent with only the NC being involved in Jab1 binding (29) (Fig. 1B) and the relative position of the CC thus not being directly influenced by this interaction partner. Thus, the altered conformation of hBrr2^T1^ in the hBrr2^T1^-hJab1^ΔCtail^ co-crystal structure is likely not induced by hJab1^ΔCtail^ binding but probably a consequence of crystal packing forces.

Notably, the same trends in unwinding rates were observed when the experiments were conducted in the presence of hJab1^ΔCtail^ (Fig. 4; Table 3). *I.e.*, in the presence of the activator, hBrr2^T1-SS-linear^ again showed a higher unwinding rate constant (2.337 ± 0.086 s^-1^) while hBrr2^T1-SS-rot+^ exhibited reduced unwinding (1.385 ± 0.055 s^-1^) compared to WT hBrr2^T1^ (1.987 ± 0.098 s^-1^; Fig. 4; Table 3).

### Altered ATP hydrolysis may contribute to altered helicase activities

To further track down the mechanism, by which the CC exerts influence on the NC depending on its relative position and orientation, we tested the consequence of stabilizing hBrr2^T1^ in different cassette configurations on ATP hydrolysis. We determined the RNA-stimulated ATPase rates of the disulfide-bridged hBrr2^T1^ variants *via* thin-layer chromatography, which again revealed significant differences compared to the WT protein, fully consistent with the trends in the unwinding activities (Fig. 5A). Thus, hBrr2^T1-SS-linear^ exhibited higher stimulated ATPase activity (7.58 ± 0.59 ATP/hBrr2^T1-SS-linear^/min), whereas the stimulated ATPase activity of hBrr2^T1-SS-rot+^ was decreased (2.18 ± 0.42 ATP/hBrr2^T1-SS-rot+^/min) compared to the WT protein (4.82 ± 0.78 ATP/hBrr2^T1^/min). The altered nucleotide binding activities of the hBrr2^T1^ variants may have contributed to the observed changes in stimulated ATPase activities.

**Figure 5.**
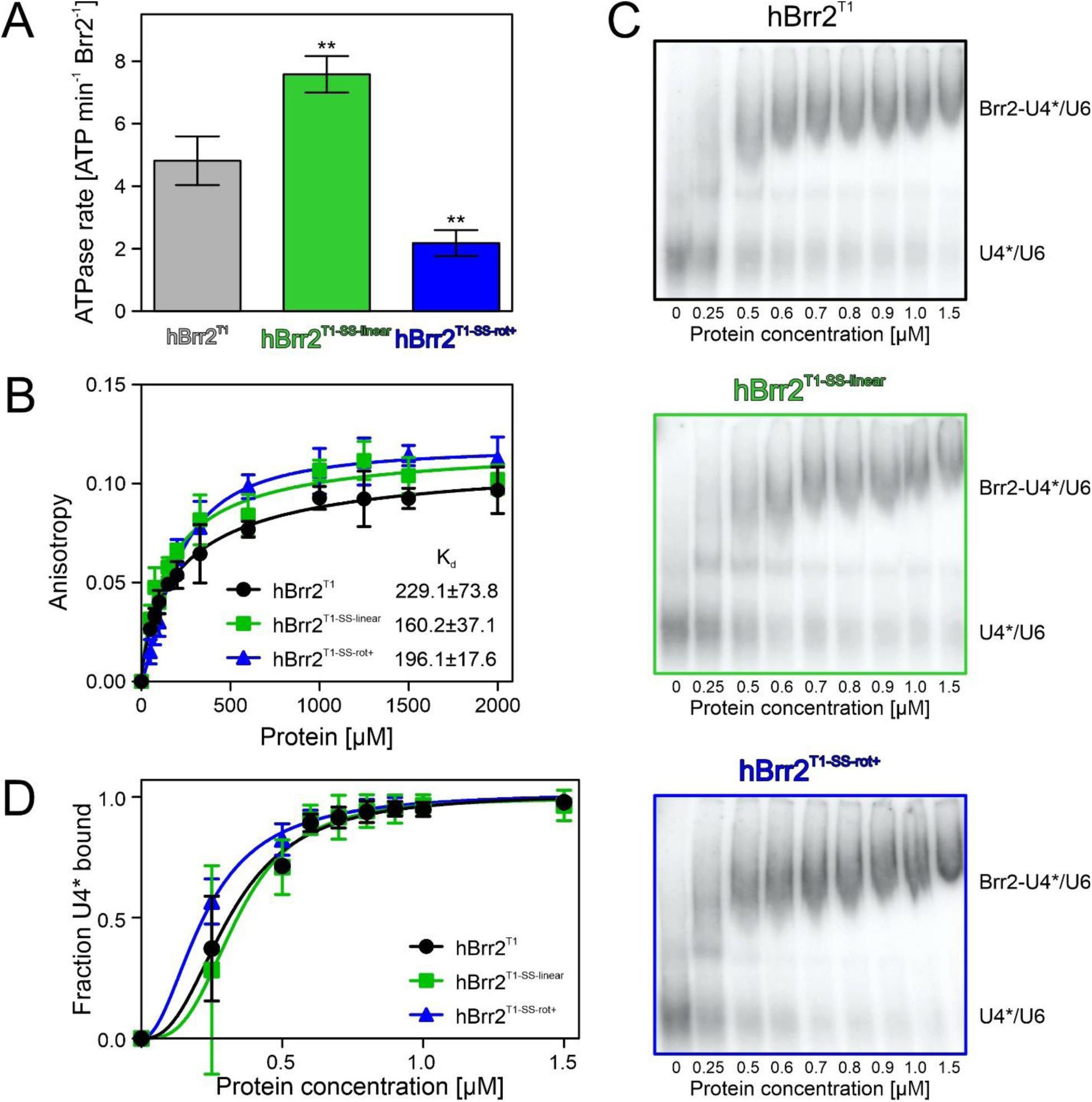
ATPase and RNA binding activities. (A) RNA-stimulated ATPase rates of the indicated hBrr2^T1^ variants (hBrr2^T1^, black; hBrr2^T1-SS-linear^, green; hBrr2^T1-SS-rot+^, blue), analyzed *via* a radioactive, thin-layer chromatography-based ATPase assay. Data points represent means ± SD of at least three independent experiments. **, p ≤ 0.01. (B) Fluorescence polarization-based analysis of RNA binding activities of the indicated hBrr2^T1^ variants (hBrr2^T1^, black; hBrr2^T1-SS-linear^, green; hBrr2^T1-SS-rot+^, blue). The indicated K_d_ values were determined by fitting the data to a Hill equation, fraction bound = A c_prot_^n^/[c_prot_^n^+Kd^n^], in which A is the maximum of bound RNA, c_prot_ is the protein concentration, K_d_ is the dissociation constant, n is the Hill coefficient. Values represent means ± SEM of at least three independent experiments. (C) Representative EMSA assays monitoring binding of the indicated hBrr2^T1^ variants (hBrr2^T1^, black; hBrr2^T1-SS-linear^, green; hBrr2^T1-SS-rot+^, blue) to U4*/U6 di-snRNAs. U4*, radioactively labeled U4 snRNA. (D) Quantification of the data shown in (C). The indicated K_d_ values were determined by fitting the quantified data to a Hill equation, fraction bound = A c_prot_^n^/[c_prot_^n^+K_d_^n^], in which A is the maximum of bound RNA, c_prot_ is the protein concentration, K_d_ is the dissociation constant, n is the Hill coefficient. Data points represent means ± SEM of at least three independent experiments.

Finally, we tested for differences in the RNA affinities of the different hBrr2^T1^ variants. Fluorescence polarization assays using a fluorescence labeled RNA duplex with a 3’-single-stranded overhang, which we had employed previously in hBrr2-RNA interaction studies (51), suggested that RNA binding was unaffected by the engineered disulfide bridges (Fig. 5B). Likewise, all proteins exhibited similar affinities to U4/U6 di-snRNAs based on electrophoretic gel mobility shift assays (EMSA; Fig. 5C,D).

## DISCUSSION

### Intra-molecular modulation in a dual-cassette RNA helicase

Peripheral domains of SF2 helicases can influence the enzymes’ subcellular localization (52), oligomerization state (53), substrate specificities (54), molecular mechanisms (55) and activities (56). The CC of Brr2 can be considered a large (about 100 kDa) and unique helicase-associated region of the N-terminal helicase unit, as it is catalytically inactive (13,34) yet influences the NC helicase from a distance (34). The functional importance of the CC is underscored by the observation that its deletion is lethal in yeast (35). However, the molecular mechanisms, by which it modulates the NC helicase are presently poorly understood.

By comparing Brr2 subunits from several crystal and cryoEM structures of isolated Brr2, small Brr2 complexes and spliceosomal complexes, we showed that the CC can adopt multiple positions and orientations relative to the NC, stabilized by different inter-cassette contacts. Locking hBrr2^T1^ in two different conformations that exhibit different relative positioning of NC and CC *via* engineered disulfide bridges, affected the helicase activity of the NC in opposite ways. We previously reported that the exchange of residues that mediate inter-cassette contacts in an isolated hBrr2^T1^ crystal structure, mutations in the inter-cassette linker and mutations in the CC nucleotide binding pocket led to reduced Brr2 helicase activity, suggesting that the CC can stimulate the activity of the NC (34). Our present observation that an engineered disulfide bridge, which stabilizes the linear cassette conformation of isolated hBrr2^T1^, leads to an increase in Brr2 helicase activity is consistent with this notion. On the other hand, a disulfide-bridged hBrr2^T1^ variant that stabilizes the enzyme in the alternative rot+ conformation, seen in the hBrr2^T1^-hJab1^ΔCtail^ co-crystal structure, led to a decreased unwinding activity. Together, these findings show that depending on its position, orientation and contacts to the NC, the CC can either activate or inhibit the NC.

We further characterized the molecular mechanism underlying this flexible regulatory influence. Our data indicate that the different NC-CC conformations do not influence RNA binding by the NC. In contrast, we observed conformation-dependent modulation of nucleotide binding and, possibly as a direct consequence, conformation-dependent modulation of stimulated ATPase rates. The trends observed in nucleotide binding and hydrolysis matched the trends seen in RNA helicase activities.

It remains to be seen if the principles uncovered here for the Brr2 RNA helicase also exist in other dual-cassette Ski2-like enzymes, such as ASCC3 and Slh1/Rqt2. In this respect, it is interesting that the CC of ASCC3 is an active DNA helicase (17), but whether the ASCC3 NC can also unwind nucleic acid duplexes and/or whether inter-cassette modulation of activity is also possible in the other direction (*i.e.* the NC influencing the activity of the CC) are open questions.

### Implications for Brr2 regulation by protein partners

In none of the presently structurally characterized splicing stages, hBrr2 adopts a conformation that corresponds to the rot+ conformation observed in the hBrr2^T1^-hJab^ΔCtail^ complex. However, not all stages of a canonical splicing reaction have so far been structurally characterized, and additional stages may be populated in certain alternative splicing events. Irrespectively, also in some of the presently structurally characterized splicing stages hBrr2 exhibits a cassette configuration distinct from the linear conformation seen in isolated hBrr2^T1^, the most extreme case represented by the rot-conformation in the B complex. As the linear conformation has a strong stimulatory effect on the NC helicase, all deviations from this conformation may lead to more or less pronounced Brr2 helicase inhibition.

Many protein interaction partners of Brr2 bind to the CC (32,33,36,37). However, the CC most likely does not simply represent a passive landing pad for other proteins, as some of the interacting proteins also influence the activity of Brr2. For example, the intrinsically disordered proteins Ntr2 (in yeast) and FBP21 (in human) bind to the CC of Brr2 and tune down its unwinding activity *in vitro* (32,33). In the respective studies, it was speculated that their binding to the CC might influence the communication between the cassettes. Based on these previous observations and results presented here, we propose a more general mechanism, wherein the CC can be used by interacting proteins to modulate the NC helicase activity from a distance, by stabilizing the CC in a helicase-promoting or in a helicase-attenuating conformation. Based on our results and our comparisons of hBrr2 structures in diverse molecular contexts, Brr2 might adopt a near continuum of different cassette conformations. The equilibrium between the conformations could be shifted by protein interaction partners binding to the CC, which might consequently gradually stimulate or inhibit the enzyme. Such protein-mediated communication between the cassettes may contribute to a retinitis pigmentosa-linked Brr2 mutation found in the CC (57).

### Implications for splicing regulation

Apart from the CC and proteins binding the CC, Brr2 activity is also regulated *via* an auto-inhibitory N-terminal region (28), *via* the Prp8 Jab1 domain that can either inhibit or activate the helicase (29-31), *via* the Prp8 RNaseH-like domain that can sequester the U4/U6 substrate (58) and *via* other proteins alter the stability of the U4/U6 substrate (9). The question emerges why Brr2 is regulated in an intricate manner on many levels and not simply shut on and off by one mechanism. We suggest that the fine tuning of Brr2 activity could be part of proofreading events or contribute to the selection of alternative splice sites.

Proofreading describes a mechanism for discriminating between suboptimal/aberrant and optimal splice substrates to maintain splicing fidelity. At least five of the spliceosome-associated NTPases/helicases, Prp5, Prp28, Prp16, Prp22, and Prp43, have been implicated in proofreading mechanisms in yeast (23), including proofreading of 5’-splice sites (59,60), of branch sites (61) and of 3’-splice sites (62). It is thought that these helicases can alternatively remodel suboptimal splicing intermediates in an off-pathway manner. After such off-pathway remodeling, often the helicase Prp43 is thought to be responsible for discarding aberrant intermediates (63). The decision or distribution between on-pathway and off-pathway remodeling may be determined by the speed at which a given helicase can remodel an intermediate along the on-pathway direction. In principle, similar mechanisms might underlie alternative splicing decisions in higher eukaryotes. Weak splice sites (analogous to suboptimal substrates) would exhibit slow splicing kinetics and thus be discarded more frequently than strong splice sites (analogous to optimal substrates) that would splice faster and thus be less prone to be discarded. Interestingly, a recent study directly linked spliceosomal helicases to alternative splicing by showing that Prp16 and Prp22 enable the selection of alternative branch sites and 3’-splice sites, respectively (24).

Some evidence suggests that also Brr2 might be involved in proofreading and alternative splicing. For example, changes in alternative splicing patterns were detected upon knockdown of core splicing factors, including Brr2 (64,65). Furthermore, two retinitis pigmentosa-linked mutations that give rise to Brr2 variants enhance the use of a cryptic splice site (66). Regulatory factors or regions, including the CC and CC-binding proteins, may modulate the kinetics of U4/U6 unwinding by Brr2, which could tune the velocity of B_act_ formation. In addition, Brr2 lacking the N-terminal region can lead to non-canonical disruption of the U4/U6•U5 tri-snRNP (into U4/U6 di-snRNP and U5 snRNP (28), and B complex spliceosomes have been shown to undergo repeated attempts at activation with intermittent release of U4 (67). Thus, Brr2 might be able to remodel the B complex in diverse ways, and thereby differentially funnel B complexes based on different pre-mRNA substrates along or off the splicing pathway. Fine tuning of Brr2 activity might then influence, to which extent a given B complex is discarded or further productively processed. Finally, unlike the other spliceosomal helicases, after formation of the B complex Brr2 remains stably associated with the spliceosome during the subsequent stages of splicing. Its helicase activity is thus in principle available to serve also as a discard factor. Although direct evidence is presently missing, tuning of the activity of Brr2 might thus even be used directly for proofreading.

## EXPERIMENTAL PROCEDURES

### Site-directed mutagenesis, cell culture and gene expression

A codon-optimized gene, encoding an N-terminal deletion construct of hBrr2 (residues 395-2129, here referred to as hBrr2 truncation 1, hBrr2^T1^), in a pFL vector, which directs the production of a fusion protein bearing an N-terminal, cleavable His_10_-tag (34), was used as a PCR template. Mutagenesis was performed with the QuikChange site-directed mutagenesis kit II (Stratagene) and successful mutagenesis was confirmed by sequencing. For production of hJab1 domain variants, we employed a DNA construct in a pFL vector that directs the production of fusion proteins with N-terminal, cleavable GST-tags (29).

The pFL constructs were transformed into *Escherichia coli* DH10 MultiBac cells and blue-white screening was used to select for colonies with successful Tn7 transposition into the baculovirus genome. The Bacmid DNA was purified and transfected into SF9 insect cells with Xtreme gene 9 DNA transfection reagent (Roche). The first virus generation, V0, was harvested and used to infect H5 insect cells to produce the second virus generation, V1. The V1 virus was used to infect H5 cells for protein production. Cells were harvested before the cell viability decreased below 90 %.

### Protein purification

All purification steps were carried out at 4 ° C. For hBrr2^T1^ variants, cell pellets were resuspended in lysis buffer (50mM HEPES-NaOH pH 7.5, 600 mM NaCl, 10 % (w/v) glycerol, 0.05 % (v/v) NP-40, 20 µg/ml DNase1, 2 mM β-mercaptoethanol, supplemented with Complete EDTA-free protease inhibitors) and sonicated for 30 min using a Sonoplus Ultrasonic Homogenizer HD 3100 (Bandelin). The lysate was centrifuged for 1 h at 21,500 rpm. The supernatant was loaded on a Histrap FF column (GE Healthcare), washed and eluted in a gradient to 250 mM imidazole. The His-tag was cleaved and the protein sample was dialyzed overnight in 40 mM HEPES-NaOH, pH 7.5, 500 mM NaCl, 10 % (w/v) glycerol, 15 mM imidazole, 2 mM β-mercaptoethanol. The cleaved protein was again loaded on a Histrap column and collected in the flow-through. The sample was then treated with RNase A (Sigma) and loaded on a HiPrep™ Heparin 16/60 column (GE Healthcare) in 25 mM TRIS-HCl, pH 8.0, 50 mM NaCl, 5 % (v/v) glycerol, 2 mM DTT, washed and eluted with a linear gradient to 750 mM NaCl. Fractions containing the protein of interest were pooled, concentrated and loaded on a HiLoad Superdex 200 16/60 column (GE Healthcare) in gel filtration buffer (10 mM TRIS-HCl, pH 7.5, 200 mM NaCl). Fractions containing the protein of interest were pooled, concentrated to 10 mg/ml, flash-frozen in liquid nitrogen and stored at -80 ° C until use. For activity assays, the protein tag was not cleaved and 20 % (v/v) glycerol were included in all buffers.

hJab1 variants were purified as described before (29). Briefly, insect cell pellets were resuspended in 50 mM TRIS-HCl, pH 8.0, 300 mM NaCl, 5 % (v/v) glycerol, 0.05 % (v/v) NP-40, 2 mM DTT, supplemented with Complete EDTA-free protease inhibitors. After sonication and centrifugation, the protein was captured on GSH beads. After washing, the protein was eluted with 10 mM reduced glutathione. The buffer was exchanged with a HiLoad Superdex 75 26/60 column to 50 mM TRIS-HCl, pH 8.0, 300 mM NaCl, 5 % (v/v) glycerol, 2 mM DTT. The GST-tag was cleaved overnight with PreScission protease and the protein was collected in the unbound fraction after adding fresh GSH beads. The protein was further purified by gel filtration in 10 mM TRIS-HCl, pH 8.0, 150 mM NaCl. Fractions containing the protein of interest were pooled, concentrated to 10 mg/ml, flash-frozen in liquid nitrogen and stored at -80 ° C until use.

For complex formation, hBrr2^T1^ variants were combined with a 1.5 molar excess of hJab1 variants and loaded on a Superdex 200 10/300 global increase column (GE Healthcare) in 20 mM TRIS-HCl, pH 8.0, 150 mM NaCl. Fractions containing the complex of interest were pooled, concentrated to 6 mg/ml, flash-frozen in liquid nitrogen and stored at -80 ° C until use.

### Crystallographic analyses

hBrr2^T1^-hJab1 complexes were crystallized in sitting drops on 24-well plates with drops containing 1 µl of protein complex solution and 1 µl of reservoir solution (0.1 M HEPES-NaOH, pH 8.0, 0.1 M MgCl_2_, 8 % PEG 3350). Crystals were cryo-protected by transfer into reservoir solution supplemented with 25 % (v/v) ethylene glycol.

Isolated hBrr2^T1^ variants were crystallized in sitting drops on 24-well plates with drops containing 1 µl of protein solution and 1 µl of reservoir solution (0.1 M Na citrate, 1.5 M Na malonate, pH 7.0). Crystals were cryo-protected in 0.1 M Na citrate, 3.0 M Na malonate, pH 7.0, 0.1 M NaCl. All crystals were flash-frozen in liquid nitrogen.

Data sets were collected on beamlines 14.1 and 14.2 of the BESSY II storage, Berlin, Germany. The data were processed with XDS (68). Structures were solved by molecular replacement with Phenix (69), using structure coordinates of isolated hBrr2^T1^ (PDB ID 4F91) and hBrr2^T1^-hJab1 complex (PDB ID 4KIT). Models were refined by automated refinement in Phenix or REFMAC5 (70) alternating with manual model building in Coot (71).

For the calculation of the relative rotation of CC with respect to NC, hBrr2 models from crystal and cryoEM structures were aligned to the crystal structure of isolated hBrr2^T1^ (PDB ID 4F91) according to Cα atoms of residues 1-1288 (NC), using the LSQ Superpose routine in Coot. Cα atoms of residues 1291-2125 (CC) of isolated hBrr2^T1^ were then aligned to the repositioned hBrr2 models from crystal and cryoEM structures. The angle α was calculated from the trace from the resulting rotation matrix 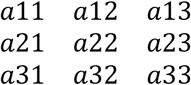 with *trace = a*11 + *a*22 + *a*33 = *n* − 2 + 2 × *cosα*, in which n was set to 3 for three-dimensional space.

### Differential scanning fluorimetry

For thermal shift analyses, hBrr2^T1^ variants were diluted to 0.2 mg/ml in 40 mM TRIS-HCl, pH 7.5, 50 mM NaCl, 0.5 mM MgCl2 with or without 2 mM DTT, and supplemented with SyproOrange Protein Gel Stain 1:500 (Sigma). After incubation on ice for 1.5 h, the protein solutions were 1:1 diluted in either 40 mM TRIS-HCl, pH 7.5, 50 mM NaCl, 0.5 mM MgCl2 with or without 2 mM DTT or in size exclusion buffer with or without 2 mM DTT. Melting curves were recorded by monitoring the fluorescence at 569 nm while heating the samples from 25 ° C to 95 ° C at 1 ° C/min.

### Mass spectrometry

Protein digestion for LC-MS analysis was conducted according to (72). Briefly, 25 µg of protein were denatured in 8 M urea, 100 mM TRIS-HCl, pH 6.5. One half of the sample was directly alkylated by addition of 2 mM N-ethylmaleimide (NEM) for 4 h at 37 ° C (non-reduced sample). The other half was first reduced by addition of 4 mM DTT for 2 h at 37 ° C, and subsequently free thiols were alkylated by addition of 2 mM NEM for 2 h at 37 ° C (reduced sample). Proteins were digested by addition of LysC protease for 4 h at 37 ° C. After a fourfold dilution in 100 mM TRIS-HCl, pH 6.5, the digestion was completed by addition of trypsin and overnight incubation at 37 ° C. The reaction was stopped by addition of 3 % (v/v) TFA, 5 % (v/v) acetonitrile, and samples were desalted by STAGE tips as described before (73).

Peptides were reconstituted in 20 μl 0.1 % (v/v) TFA, 5 % (v/v) acetonitrile in water and 2 µl were analyzed by a reverse-phase capillary nano liquid chromatography system (Ultimate 3000, Thermo Scientific) connected to an Orbitrap Velos mass spectrometer (Thermo Scientific). Samples were desalted on a trap column (PepMap100 C18, 3 μm, 100 Å, 75 μm i.d. × 2 cm; Thermo Scientific) using a mobile phase of 0.05 % (v/v) TFA, 2 % (v/v) acetonitrile in water. LC separations were performed on a capillary column (Acclaim PepMap100 C18, 2 μm, 100 Å, 75 μm i.d. × 25 cm; Thermo Scientific) at an eluent flow rate of 300 nl/min. Mobile phase A contained 0.1 % (v/v) formic acid in water, and mobile phase B contained 0.1 % (v/v) formic acid in acetonitrile. The column was pre-equilibrated with 3 % mobile phase B followed by a linear increase to 50 % mobile phase B over 50 min. Mass spectra were acquired in a data-dependent mode, utilizing a single MS survey scan (R = 30,000) in the Orbitrap, followed by up to 20 HCD scans (R = 7,500), using a normalized collision energy of 35 excluding +1 precursor ions. Extracted ion chromatograms (XIC) and spectra were analyzed by Thermo Scientific Xcalibur 2.2.

### U4*/U6 unwinding assays

U4*/U6 snRNA duplex (U4* - U4 radioactively labeled) was prepared as described previously (9). Briefly, full-length yeast U4* snRNA was annealed with full-length U6 snRNA in 20 mM TRIS-HCl, pH 7.5, 50 mM NaCl, 18 mM MgCl_2_. Annealed duplex was separated from single strands by a 6 % native PAGE at 200 V for 1 h. The duplex regions were extracted from the gel in 20 mM TRIS-HCl, pH 8.0, 300 mM NaCl, 10 mM EDTA, 0.5 % (v/v) SDS, further purified by phenol-chloroform extraction and precipitated with LiCl and isopropanol. The RNA was washed with 70 % ethanol and dissolved in 40 mM TRIS-HCl, pH 7.5, 100 mM NaCl.

For unwinding assays, 200 nM hBrr2^T1^ variants with or without 500 nM hJab1^ΔCtail^ were incubated for 3 min at 30 ° C with 0.65 nM U4*/U6 duplex in unwinding buffer, supplemented with 8% (v/v) glycerol, 15 ng/µl acetylated BSA, 1 U/µl RNasin (MoloX), with or without 2 mM DTT. The unwinding reactions were started by addition of 1.7 mM ATP/MgCl_2_. 10 µl samples were taken at the indicated time points and the reactions were stopped by mixing with 40 mM TRIS-HCl, pH 7.4, 50 mM NaCl, 25 mM EDTA, 1 % (w/v) SDS, 10 % (v/v) glycerol, 0.05 % (w/v) xylene cyanol, 0.05 % (w/v) bromophenol blue. The samples were loaded on a 6 % native PAGE and separated at 200 V, 4 ° C for 1 h. The gels were transferred to a filter paper, visualized by autoradiography and bands were quantified with the ImageQuant software. The data of three independent replicates were fitted to the equation y=A{1-exp[-*k*_*u*_t]}, in which y is the fraction U4*/U6 unwound, A is the amplitude of the reaction, *k*_*u*_ is the apparent rate constant of unwinding and t is time, by using GraphPad Prism.

### ATPase assays

ATP hydrolysis reactions were started by addition of 1 mM ATP/MgCl2, supplemented with a small amount of α-^P32^ ATP, to 100 nM hBrr2^T1^ variants and 1 µM U4/U6 duplex in unwinding buffer. Samples were incubated for 30 min at 30 ° C, reactions were stopped by addition of 50 mM EDTA, and samples were transferred to a TCL PEI cellulose F plate (Merck). Thin-layer chromatography was run with 20 % (v/v) EtOH, 6 % (v/v) acetic acid, 0.5 M LiCl. The plate was dried and ATP/ADP signals were visualized by autoradiography. Radioactive spots were quantified with the ImageQuant software, and the number of hydrolyzed ATP molecules per hBrr2^T1^ variant molecule per minute was calculated based on three independent replicates.

### Fluorescence polarization assays

10 nM FAM-labeled RNA duplex bearing a 3’-single stranded (ss) overhang (5’-**CGCGUGCUGGUC**GAAAUUUAAUUAUAA ACCAGACCGUC-3’/5’-**GACCAGCACGCG**-3’-[5-FAM]); regions forming the duplex in bold; IBA) were incubated with increasing concentrations of hBrr2^T1^ variants (0, 50, 75, 100, 150, 200, 500, 1000, 1250, 1500, 2000 nM) in 40 mM HEPES-NaOH, pH 7.5, 50 mM NaCl, 0.5 mM MgCl_2_, 8 % (v/v) glycerol, 10 nM acetylated BSA at room temperature. Samples were measured in a Victor V3 1420 plate reader. The fluorescence polarization values for three independent replicates were plotted with GraphPad Prism. K_d_’s were determined by fitting the data to a Hill equation, fraction bound = A c _prot_ ^n^/[c _prot_ ^n^+K _d_ ^n^], in which A is the maximum of bound RNA, c_prot_ is the protein concentration, Kd is the dissociation constant, n is the Hill coefficient.

### Electrophoretic mobility shift assays

Increasing concentrations of hBrr2^T1^ variants (0, 0.25, 0.5, 0.6, 0.7, 0.8, 0.9, 1.0, 1.5, 2.0, 3.0 µM) were incubated with 1 nM radioactive U4*/U6 duplex RNA for 3 min at 30 ° C in unwinding buffer, supplemented with 0.25 mg/ml yeast tRNA. The samples were mixed with 40 mM TRIS-HCl, pH 7.4, 50 mM NaCl, 25 mM EDTA, 10 % (v/v) glycerol, 0.05 % (w/v) xylene cyanol, 0.05 % (w/v) bromophenol blue and run on a 4 % native PAGE at 170 V, 4 ° C for 2-3 h. The gels were transferred to filter papers and visualized by autoradiography. Quantification of three independent replicates was done with ImageQuant software and plotted with GraphPad Prism. K_d_’s were determined by fitting the quantified data to a Hill equation, fraction bound = A c_prot_^n^/[c_prot_^n^+K _d_ ^n^], in which A is the maximum of bound RNA, c_prot_ is the protein concentration, K_d_ is the dissociation constant, n is the Hill coefficient.

## Acknowledgements

We thank Eva Absmeier, Freie Universität Berlin, for helpful discussions. We accessed beamlines of the BESSY II storage ring (Berlin, Germany) *via* the Joint Berlin MX-Laboratory (https://www.helmholtz-berlin.de/forschung/oe/np/gmx/joint-mx-lab) sponsored by Helmholtz Zentrum Berlin für Materialien und Energie, Freie Universität Berlin, Humboldt-Universität zu Berlin, Max-Delbrück Centrum for Molecular Medicine, Leibniz-Institut für Molekulare Pharmakologie and Charité – Universitätsmedizin Berlin. For mass spectrometry (B.K. and C.W.), we acknowledge the assistance of the Core Facility BioSupraMol supported by the Deutsche Forschungsgemeinschaft (DFG). This work was funded by grant TRR186/A15-1 from the Deutsche Forschungsgemeinschaft to MCW and by a Dahlem International Network PostDoc Fellowship from Freie Universität Berlin to KFS.

## Footnotes

This work was funded by grant TRR186/A15-1 from the Deutsche Forschungsgemeinschaft to MCW and by a Dahlem International Network PostDoc Fellowship from Freie Universität Berlin to KFS.

The abbreviations used are: BESSY II, Berliner Elektronenspeicherring-Gesellschaft für Synchrotronstrahlung II, CC, C-terminal cassette; cryoEM, cryo-electron microscopy; DSF, differential scanning fluorimetry; DTT, dithiothreitol; EMSA, electrophoretic gel moibility shift assay; h, human; hBrr2^T1-SS-linear^, hBrr2^T1-M641C/A1582C^; hBrr2^T1-SS-rot+^, hBrr2^T1-D534C/N1866C^; HB, helical bundle domain; HEPES, 4-(2-hydroxyethyl)-1-piperazineethanesulfonic acid; HLH, helix-loop-helix domain; IG, immunoglobulin-like domain; MS, mass spectrometry; NC, N-terminal cassette; NEM, N-ethylmaleimide; NTPase, nucleic acid-dependent nucleotide tri-phosphatase; pre-mRNA, precursor messenger RNA; rmsd, root-mean-square deviation; RNP, ribonucleoprotein complex; SD, standard deviation; SEC, size exclusion chromatography; SEM, standard error of the mean; sn, small nuclear; ss, single-stranded; T1, truncation 1; TFA, trifluoroacetic acid; TRIS, tris(hydroxymethyl)aminomethane; v/v, volume/volume; WH, winged-helix domain; WT, wild type; w/v, weight/volume.

## SUPPORTING INFORMATION

### SUPPORTING FIGURES

**Figure S1.**
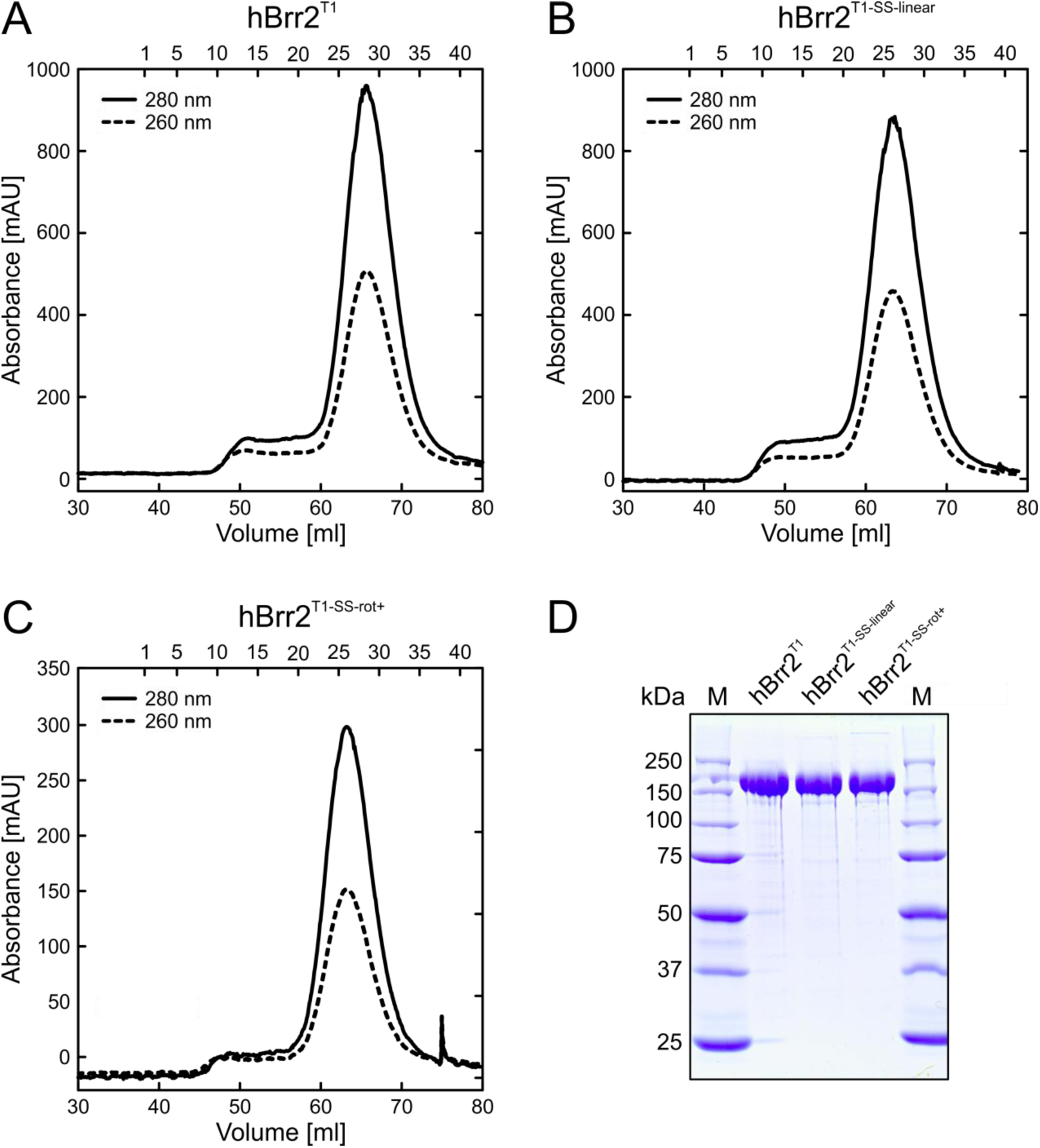
Sample quality. (A-C). Final SEC runs of the indicated, purified hBrr2^T1^ variants. Solid lines, absorbance at 280 nm; dashed lines, absorbance at 260 nm. (D) SDS-PAGE analysis of the purified hBrr2^T1^, hBrr2^T1-SS-linear^ and hBrr2^T1-SS-rot+^ samples in non-reducing loading buffer, showing the lack of formation of non-specific inter-molecular disulfide bridges.

**Figure S2.**
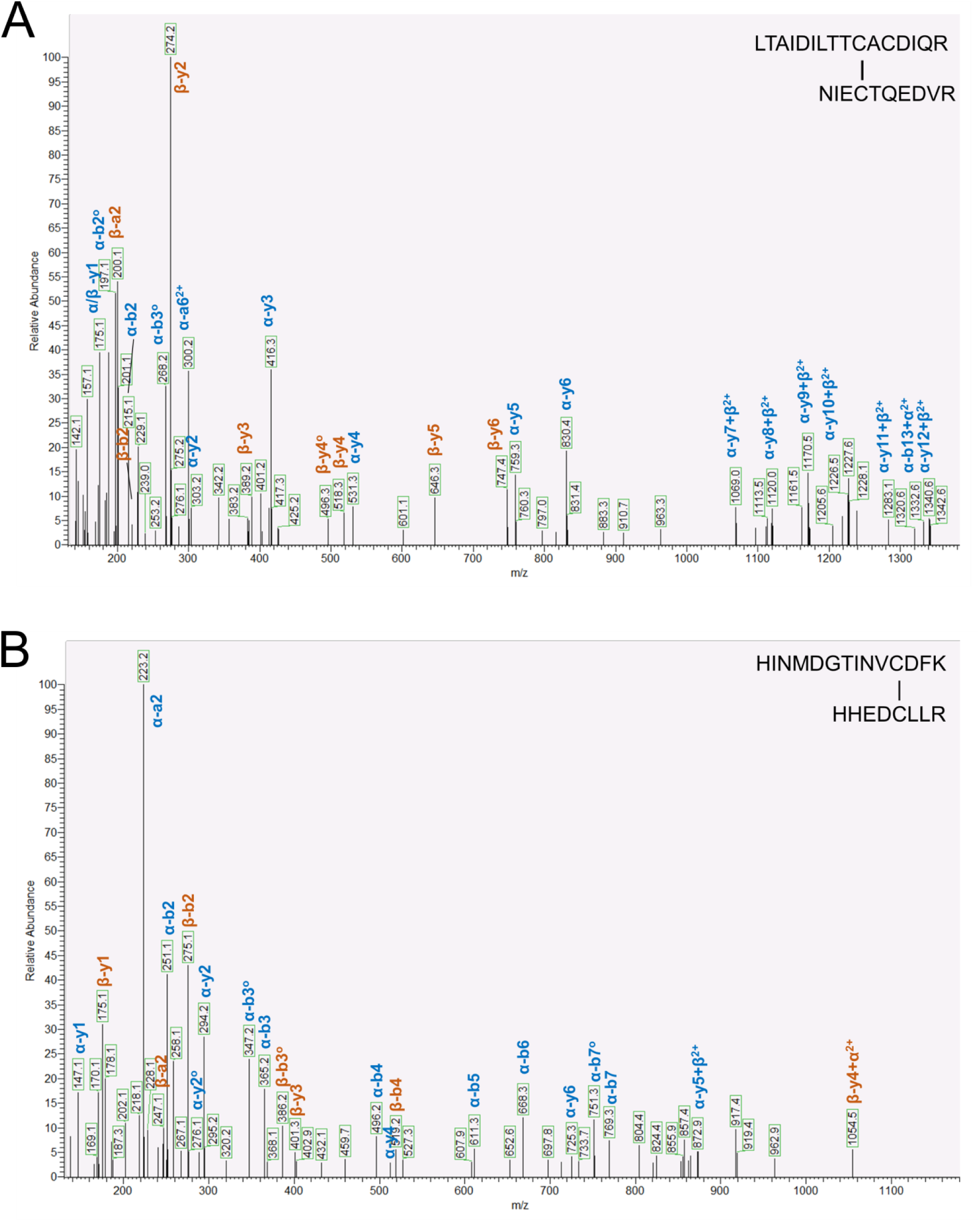
LC-MS/MS analyses. (A) LC-MS/MS analysis of in-solution digested hBrr2^T1-SS-linear^. MS2 spectrum of the quadruply charged signal at m/z 770.37, corresponding to the disulfide-linked peptides LTAIDILTTCACDIQR (α) and NIECTQEDVR (β). The presence of NEM-modified (Δm=+125 Da) fragment ions α-y5 and α-y6 of the peptide LTAIDILTTCACDIQR indicates that the engineered cysteine at position 12 is not involved in disulfide-bridge formation. Therefore, the cysteine at position 10 forms thedisulfide bridge. (B) LC-MS/MS analysis of in-solution digested hBrr2^T1-SS-rot+^. MS2 spectrum of the quadruply charged signal at m/z 657.30, corresponding to the disulfide-linked peptides HINMDGTINVCDFK (α) and HHEDCLLR (β).

